# Transposable element RNA dysregulation in mutant KRAS(G12C) 3D lung cancer spheroids

**DOI:** 10.1101/2023.02.27.530369

**Authors:** David Carrillo, Roman E. Reggiardo, John Lim, Gary Mantalas, Vikas Peddu, Daniel H. Kim

## Abstract

Mutant KRAS regulates transposable element (TE) RNA and interferon-stimulated gene (ISG) expression, but it remains unclear whether diverse mutations in KRAS affect different TE RNAs throughout the genome. We analyzed the transcriptomes of 3D human lung cancer spheroids that harbor KRAS(G12C) mutations to determine the landscape of TE RNAs regulated by mutant KRAS(G12C). We found that KRAS(G12C) signaling is required for the expression of LINE- and LTR-derived TE RNAs that are distinct from TE RNAs previously shown to be regulated by mutant KRAS(G12D) or KRAS(G12V). Moreover, KRAS(G12C) inhibition specifically upregulates SINE-derived TE RNAs from the youngest Alu subfamily AluY. Our results reveal that TE RNA dysregulation in KRAS-driven lung cancer cells is mutation-dependent, while also highlighting a subset of young, Alu-derived TE RNAs that are coordinately activated with innate immunity genes upon KRAS(G12C) inhibition.

## INTRODUCTION

Transposable element (TE) RNAs are recurrently dysregulated in the context of cancer ^1^. In mutant KRAS lung cancer cells, KRAB zinc-finger (KZNF) genes are broadly downregulated, leading to the aberrant upregulation of TE RNAs derived from LINE, SINE, and LTR elements ^2,3^. In addition to TE RNAs, long noncoding RNAs (lncRNAs) are also coordinately regulated with RAS signaling genes ^4^, and their expression patterns are similarly altered in many cancers ^5-8^. For TE RNAs, their upregulation in cancer induces a state of viral mimicry, leading to the intrinsic activation of innate immunity genes such as interferon-stimulated genes (ISGs) ^3,9,10^. In particular, the Alu family of SINEs are a predominant source of immunogenic TE RNAs ^11^, which are induced by epigenetic changes caused by DNA methyltransferase inhibitors (DNMTi) or mutant KRAS-mediated KZNF inhibition ^3,9,10^.

To investigate how a common mutation in KRAS affects the TE RNA landscape in lung cancer cells, we characterized the transcriptomes of 3D lung cancer spheroids that harbor KRAS(G12C) mutations in the presence or absence of mutant KRAS(G12C) inhibitor ^12^. We show that KRAS(G12C) signaling is required for the expression of LINE- and LTR-derived TE RNAs, while KRAS(G12C) inhibition specifically upregulates both SINE-derived TE RNAs and a subset of interferon (IFN)-related genes. Our findings reveal the complex interplay between mutant KRAS signaling and TE dysregulation in lung cancer cells, where a defined set of young AluY elements are upregulated in a mutation-dependent manner.

## RESULTS

### KRAS(G12C) inhibition alters the coding and noncoding transcriptome

To determine how oncogenic KRAS(G12C) signaling regulates the coding and noncoding transcriptome, we performed RNA sequencing (RNA-seq) on 3D mutant KRAS(G12C) lung cancer spheroids. For RNA-seq, we used a full-length protocol with 5’ unique molecular identifiers (UMIs) to enable precise RNA counting while mitigating PCR amplification biases ^13^ (Figure 1A). We compared the transcriptomes of H358 lung cancer spheroids treated with the KRAS(G12C) inhibitor ARS-1620 (ARS) to control spheroids (DMSO-treated) (Figure S1A) and saw that ARS treatment substantially reduced the levels of phosphorylated ERK (p-ERK) (Figure S1B), indicating that ARS treatment was inhibiting downstream KRAS(G12C) signaling. The suppression of KRAS(G12C) signaling by ARS treatment was also evidenced by reductions in spheroid size and cell viability (Figure S1C and S1D).

**Figure 1.**
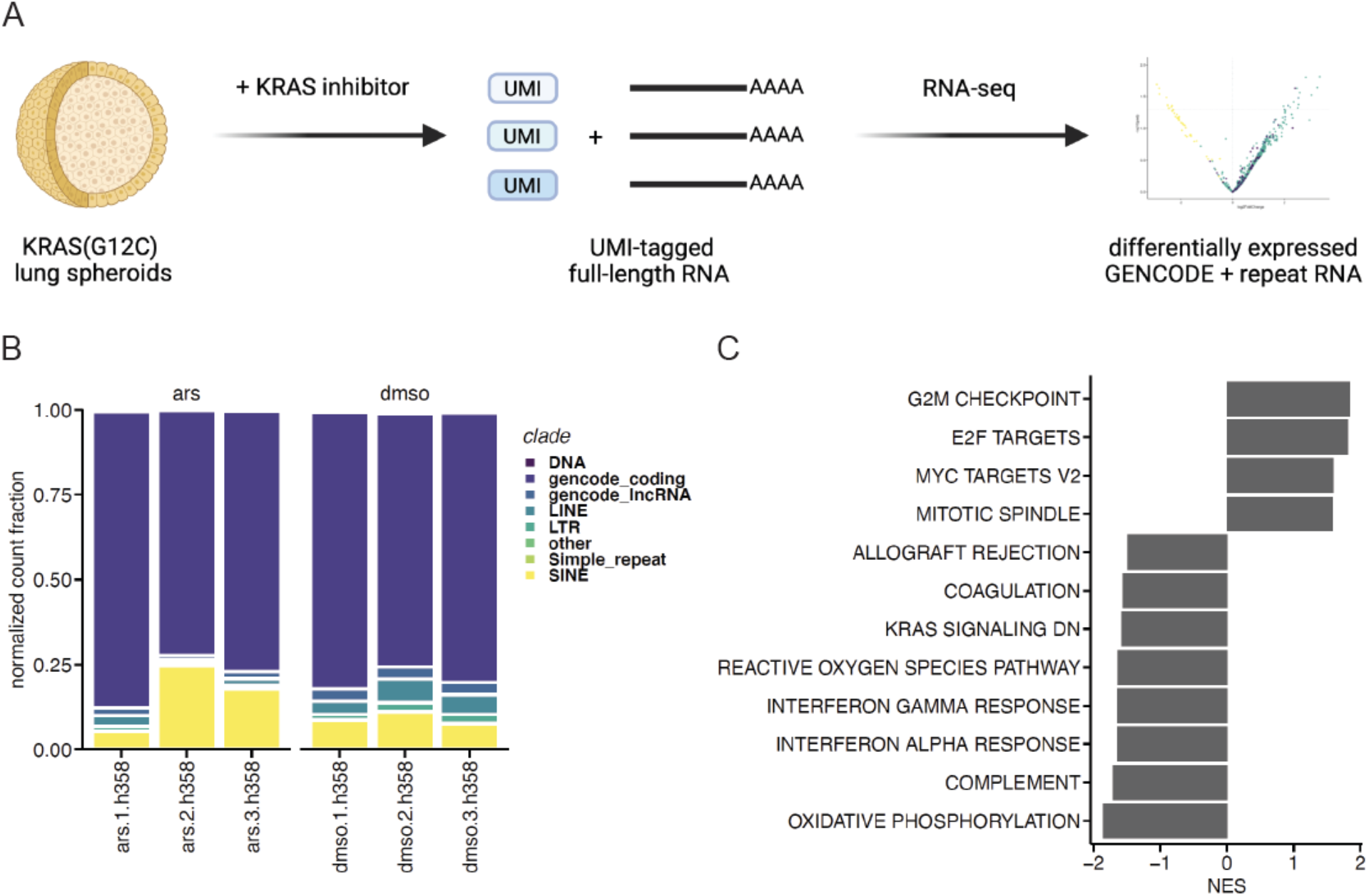
KRAS(G12C) inhibition alters the coding and noncoding transcriptome. **A**. Experimental schematic. **B**. Distribution of counts assigned to GENCODE coding, lncRNA, and TE/repeat superfamilies in ARS-treated (ars) or DMSO-treated (dmso) lung cancer spheroid RNA-seq libraries, where each column represents a biological replicate. **C**. Significant Gene Set Enrichment Analysis results observed in DMSO-treated (right, positive NES) or ARS-treated (left, negative NES) lung cancer spheroids using differentially expressed genes ranked by normalized enrichment score (NES).

At the RNA level, we assessed the relative abundances of different biotypes and TE superfamilies in ARS- or DMSO-treated 3D lung cancer spheroids, which revealed dynamic changes in TE RNA composition upon KRAS(G12C) inhibition (Figure 1B). While we detected only 32% of GENCODE-annotated protein-coding genes and 15% of lncRNA genes, 92-96% of TE superfamilies (92% LTR, 95% SINE, 96% LINE) were represented in our UMI-tagged RNA-seq data, revealing broad dysregulation of TE RNAs in the KRAS(G12C)-driven transcriptome. Up to a quarter of all detected RNA molecules in ARS-treated lung cancer spheroids were derived from TEs, indicating that TE RNAs represent a sizable portion of the transcriptional output of mutant KRAS(G12C) lung cancer cells.

We next determined the significantly differentially expressed genes between ARS- and DMSO-treated 3D lung cancer spheroids to identify biological processes that were regulated by oncogenic KRAS(G12C) signaling. Lung cancer spheroids with intact KRAS(G12C) signaling were significantly enriched for genes involved in G2M checkpoint, E2F targets, MYC targets, and mitotic spindle (Figure 1C). Upon KRAS(G12C) inhibition, however, lung cancer spheroids expressed significantly higher levels of genes involved in oxidation phosphorylation, complement, and the IFN alpha and gamma responses (Figure 1C), providing further supporting evidence for the involvement of KRAS signaling in the regulation of IFN-related genes ^2,3^.

### KRAS(G12C) inhibition coordinately induces ISGs and young AluY elements

To further elucidate the intrinsic upregulation of ISGs upon KRAS(G12C) inhibition, we identified which genes and TEs were significantly differentially expressed in each gene set and TE superfamily, respectively. Across both IFN alpha and IFN gamma response genes, the most strongly induced gene upon KRAS(G12C) inhibition in lung cancer spheroids was RTP4 (Figure 2A), a receptor transporter protein that negatively regulates TBK1 signaling ^14^. Additionally, the MHC class I complex gene beta-2-microglobulin (B2M), which is recurrently inactivated in lung cancer ^15^, was also significantly upregulated in both IFN-related gene sets in ARS-treated lung cancer spheroids (Figure 2A).

**Figure 2.**
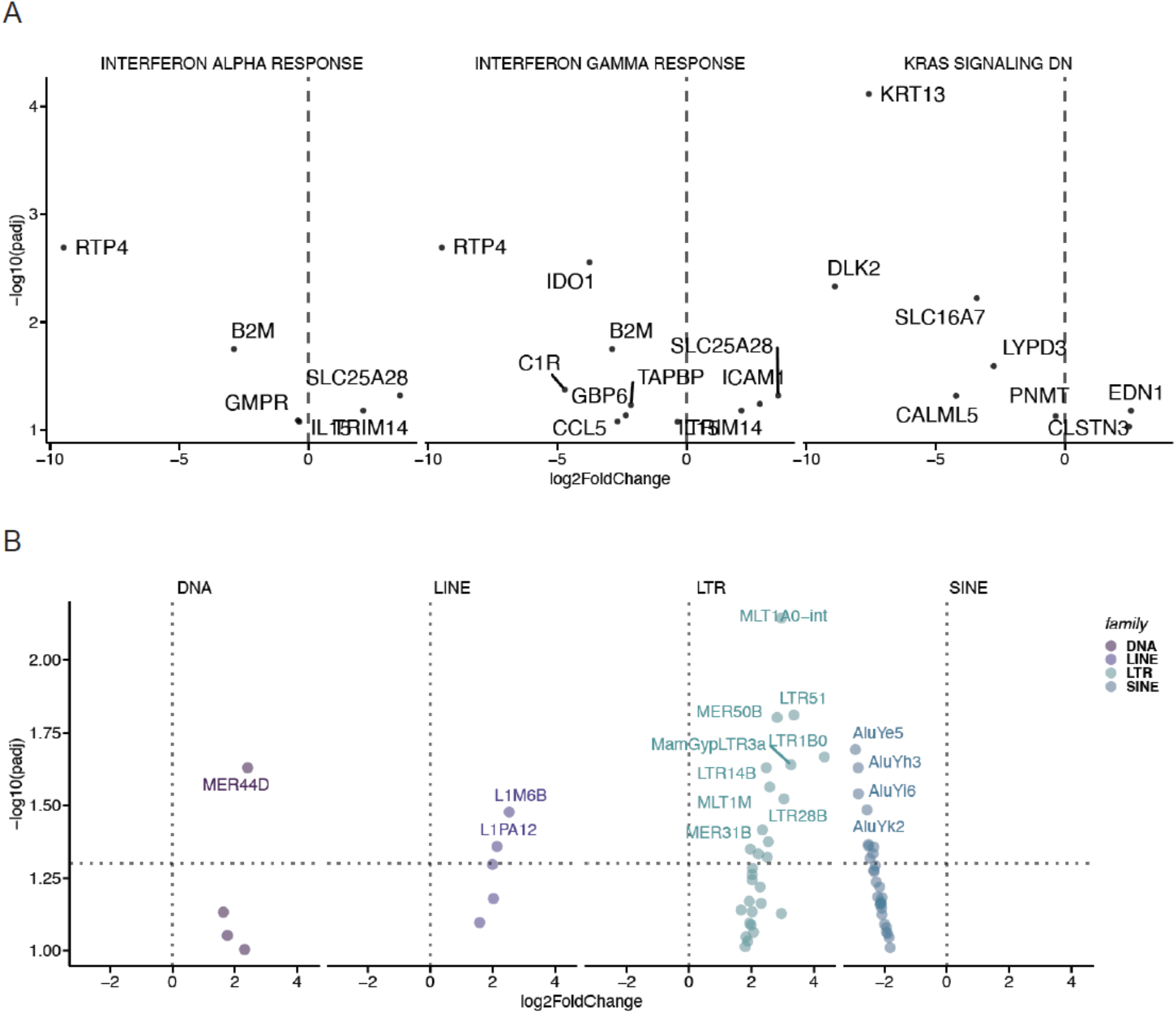
KRAS(G12C) inhibition coordinately induces ISGs and young AluY elements. **A**. Volcano plots depicting significant differential expression observed in key gene sets between DMSO-treated (right, positive fold-change) or ARS-treated (left, negative fold-change) lung cancer spheroids. **B**. Volcano plots depicting significant differential expression observed in TE superfamilies between DMSO-treated (right, positive fold-change) or ARS-treated (left, negative fold-change) lung cancer spheroids.

Given the direct role for oncogenic KRAS signaling in TE RNA regulation ^3^, we investigated which subfamilies of TE RNAs were dependent on mutant KRAS(G12C). In lung cancer spheroids with intact KRAS(G12C) signaling, we found that the LINE subfamilies L1M6B and L1PA12 were highly expressed and dependent on KRAS(G12C), as evidenced by their downregulation in lung cancer spheroids treated with ARS (Figure 2B). Moreover, while only a single DNA subfamily MER44D was dependent on KRAS(G12C) signaling, over a dozen LTR subfamilies were regulated by mutant KRAS(G12C), including MLT1A0-int, LTR51, MER50B, and LTR1B0 (Figure 2B). In contrast, SINE subfamilies were all significantly upregulated upon KRAS(G12C) inhibition and coordinately induced with specific ISG genes (Figure 2A) in lung cancer spheroids. These SINE TE RNAs were all derived from the AluY subfamily (Figure 2B), indicating that KRAS(G12C) inhibition dysregulates a specific subset of young AluY elements.

### KRAS(G12C) inhibition downregulates long noncoding RNAs

To further elucidate the effects of KRAS(G12C) inhibition on the transcriptome, we examined all significantly differentially expressed lncRNAs in lung cancer spheroids treated with ARS or DMSO control. We found that a large number of lncRNAs were dependent on KRAS(G12C) signaling for their expression, as many of these lncRNAs were significantly downregulated upon KRAS(G12C) inhibition (Figure 3A). Three of these downregulated lncRNAs, AC114546.3, NCMAP-DT, and AC073575.2, have no known functions but overlap in an antisense orientation to the coding genes ZNF770, RCAN3, and ERP29, respectively. As a class of noncoding RNA, we observed broad downregulation of lncRNAs upon ARS treatment in lung cancer spheroids (Figure 3B), indicating that many lncRNAs are dependent on KRAS(G12C) signaling for their expression.

**Figure 3.**
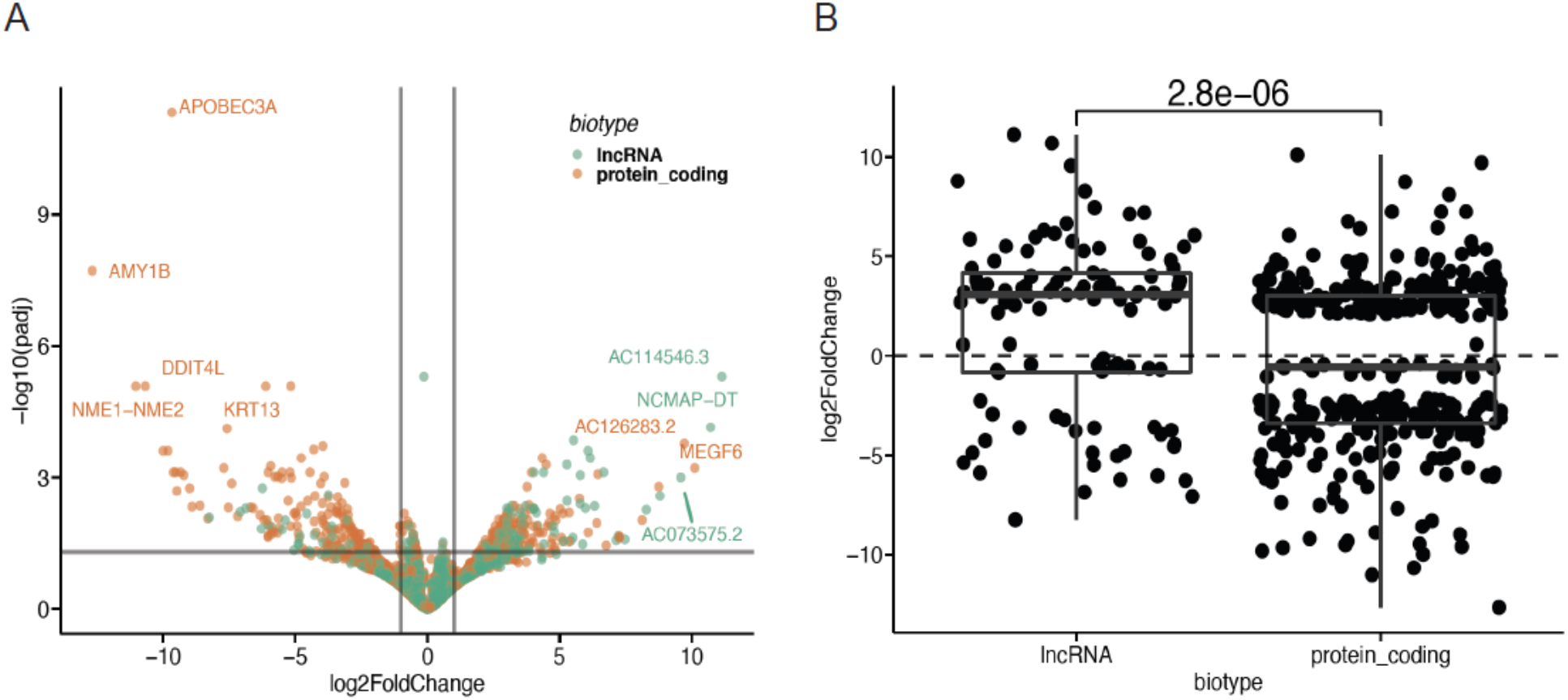
KRAS(G12C) inhibition downregulates long noncoding RNAs. **A**. Volcano plot of significant differential expression of GENCODE protein-coding RNAs and lncRNAs between DMSO-treated (right, positive fold-change) or ARS-treated (left, negative fold-change) lung cancer spheroids. **B**. Box plot of significant differential expression of GENCODE protein-coding RNAs and lncRNAs between DMSO-treated (top, positive fold-change) or ARS-treated (bottom, negative fold-change) lung cancer spheroids (Wilcoxon).

## DISCUSSION

Here we show that oncogenic KRAS(G12C) signaling is required for the expression of specific TE superfamilies, namely LINE and LTR elements, as well as a subset of lncRNAs, further demonstrating how RAS signaling regulates the noncoding transcriptome ^2-4^. We also used a 3D lung cancer spheroid model for our KRAS(G12C) inhibitor experiments since 3D models have been shown to more faithfully recapitulate the in vivo drug response when compared to 2D culture models ^16^. Furthermore, our application of a UMI-based full-length RNA-seq technique ^13^ allowed us to more accurately capture TE RNA composition and dynamics in our lung cancer spheroids by removing PCR duplicates in our RNA-seq data.

Our findings are consistent with previous studies of KRAS(G12C) inhibition, where IFN alpha and gamma response genes were upregulated in ARS-treated H358 lung cancer cells ^17^. Based on the known immunogenic properties of Alu-derived RNAs ^11^, our results suggest that the specific upregulation of young AluY elements upon KRAS(G12C) inhibition is at least in part responsible for the strong upregulation of ISGs in ARS-treated lung cancer spheroids. Notably, the significant enrichment of IFN-related genes in response to KRAS(G12C) inhibitor treatment does not include the further upregulation of RNA sensor ISGs such as MDA-5, RIG-I, or PKR, which initially become upregulated in lung cells in response to oncogenic KRAS signaling ^2,3^.

We have previously shown that mutant KRAS signaling alone is sufficient to induce TE RNA upregulation in human lung cells that have been transformed in vitro ^2,3^, and our results described here extend these observations to lung cancer cells with a different activating KRAS mutation. KRAS(G12D) or KRAS(G12V) mutations both induce the significant upregulation of the LTR12C subfamily in transformed lung cells ^3^, but we did not see significant enrichment of LTR12C-derived TE RNAs in our KRAS(G12C) lung cancer spheroids. Instead, we saw upregulation of LTR51, LTR1B0, LTR14B, and LTR28B, suggesting that diverse gain-of-function mutations in KRAS regulate different aspects of the TE RNA transcriptome.

Our work provides a comprehensive assessment of how the noncoding/TE RNA transcriptome dynamically responds to KRAS(G12C) inhibition. Future studies may provide new insights into the potential roles of noncoding/TE RNAs in mechanisms of KRAS(G12C) inhibitor resistance. Furthermore, TE RNAs that are secreted from cancer cells upon KRAS inhibition may serve as extracellular RNA biomarkers ^2,3,5,18,19^ of response and/or resistance to KRAS inhibitor therapies ^20^.

## MATERIALS AND METHODS

### Cell lines

H358 lung cancer cell lines containing the KRAS(G12C) mutation were cultured in RPMI 1640 medium (Invitrogen) supplemented with 10% fetal bovine serum (Sigma) at 37°C, 5% CO2 in a humidified incubator. All cell lines tested negative for mycoplasma. Cell lines were purchased from American Type Culture Collection (ATCC).

### Cell viability assays

For spheroid viability assays, 10,000 cells/well were seeded in low adhesion round bottom 96-well plates and incubated at 37°C, 5% CO2 for 24 hours. Then serially diluted ARS-1620 or DMSO were added to the cells, and plates were incubated in standard culture conditions for 72 hours, with fresh ARS and DMSO media being replaced daily. Cell viability was measured using a Cell Titer-Glo® Luminescent Cell Viability Assay kit (Promega) according to the manufacturer protocol. The luminescence signal of ARS-treated samples was normalized to DMSO control. Luminescence was measured on a SpectraMax iD3 molecular device.

### RNA isolation

Total bulk RNA was isolated from approximately 100 H358 spheroids (per condition) using Quick-RNA Mini-Prep kit (Zymogen) according to the manufacturer protocol. RNA was quantified via NanoDrop-8000 Spectrophotometer.

### RNA-seq library preparation

An adapted Smart-seq3 protocol ^13^ was used to generate RNA-seq libraries from total RNA to count and assess full-length RNA molecules. Briefly, 10ng of total RNA was reverse transcribed using a barcoded oligoDT primer (125nM) followed by template switching with a barcoded template switch oligo (125nM). These oligo sequences served as primers for PCR amplification. The Nextera HT kit (Illumina) was used to convert cDNA libraries into sequencing libraries with the addition of a UMI-specific primer to amplify the cDNA ends containing molecular barcodes as described in the Smart-seq3 protocol. cDNA and library quality were assessed using an Agilent bioanalyzer DNA high sensitivity chip and quantified using the high sensitivity DNA assay on the Qubit 3.0.

### Western blot

Approximately 100 H358 spheroids (per condition) were isolated following ARS or DMSO treatment. Spheroids were then incubated on ice in RIPA buffer supplemented with protease inhibitor for 15 minutes. Lysates were then spun at 10,000 RCF for 10 minutes. Supernatant was then transferred to a new tube for subsequent SDS-PAGE sample prep in Laemmli buffer, boiled for 5 minutes at 95 C for a final concentration of 1mg/ml. SDS-PAGE was performed to separate protein by size followed by subsequent transfer to a PVDF membrane. Membranes were incubated with primary p-ERK (CST) and HSP90 (CST) antibodies overnight at 4 C. Secondary antibodies (Abcam) were then incubated on 3 times TBST washed membranes in blocking buffer for subsequent imaging.

### UMI deduplication

Paired end illumina reads were adapter trimmed using FastP ^21^ with default settings. UMIs were extracted from the read and moved to the readname using umi_tools_extract from the UMI-tools package ^22^ with the barcode pattern set to “NNNNNNNN”. UMI-removed reads were aligned against HG38 using the STAR aligner with the GENCODE v38 annotation set. Aligned reads were deduplicated using “UMI-tools dedup” with default settings.

### RNA-seq analysis

All *fastq* files were trimmed with *Trimmomatic 2 (0*.*38)* ^23^ and resulting trimmed files were assessed with *FastQC* ^24^ and then processed with the following analytical pipeline: *Salmon (1*.*3*.*0):* pseudoalignment of RNA-seq reads performed with *Salmon* ^*25*^ using the following arguments:

--validateMappings –gcBias --seqBias --recoverOrphans --rangeFactorizationBins 4 using an index created from the *GENCODE* version 35 transcriptome fasta file using decoy sequences to enable selective alignment. An additional, TE-aware index was created in a similar fashion but supplemented with sequences generated from the UCSC Repeat Masker track.

*DESeq2 (1*.*32*.*0): Salmon* output was imported into a DESeq object using *tximport* ^26^ and differential expression analysis was performed with standard arguments ^27^. All results were filtered to have padj < 0.05. Where count data was used, it was normalized across samples using DESeq.

### Gene set enrichment analysis

Differentially expressed genes were ranked by the shrunken log2FoldChange values generated by *DESeq2*. Gene sets were acquired using the *R* package *msigdbr (7*.*4*.*1)* and filtered to only contain gene sets with ‘Hallmark’ status.

The *R* package *fgsea (1*.*18*.*0)* was used to generate Gene Set Enrichment estimates which were filtered to results with adjusted pvalues < 0.05.

## ACKNOWLEDGEMENTS

We thank members of the Kim Lab for helpful discussions. This work was supported by funds from the Baskin School of Engineering (to D.H.K). D.C. was supported by a Predoctoral Fellowship Award from the Tobacco-Related Disease Research Program (T30DT0997), R.E.R. was supported by an F99/K00 NIDDK KUH Predoctoral to Postdoctoral Fellow Transition Award from the National Institutes of Health (1F99DK131504-01), and V.P. was supported by a Predoctoral Fellowship Award from the Tobacco-Related Disease Research Program (T32DT4904).

## AUTHOR CONTRIBUTIONS

D.H.K. conceptualized research, D.C. and D.H.K. designed research, D.C., J.L., and G.M. performed experiments, R.E.R. and V.P. analyzed data, and D.C. and D.H.K. wrote the paper with input from the authors.

## SUPPLEMENTARY FIGURE

**Figure S1.**
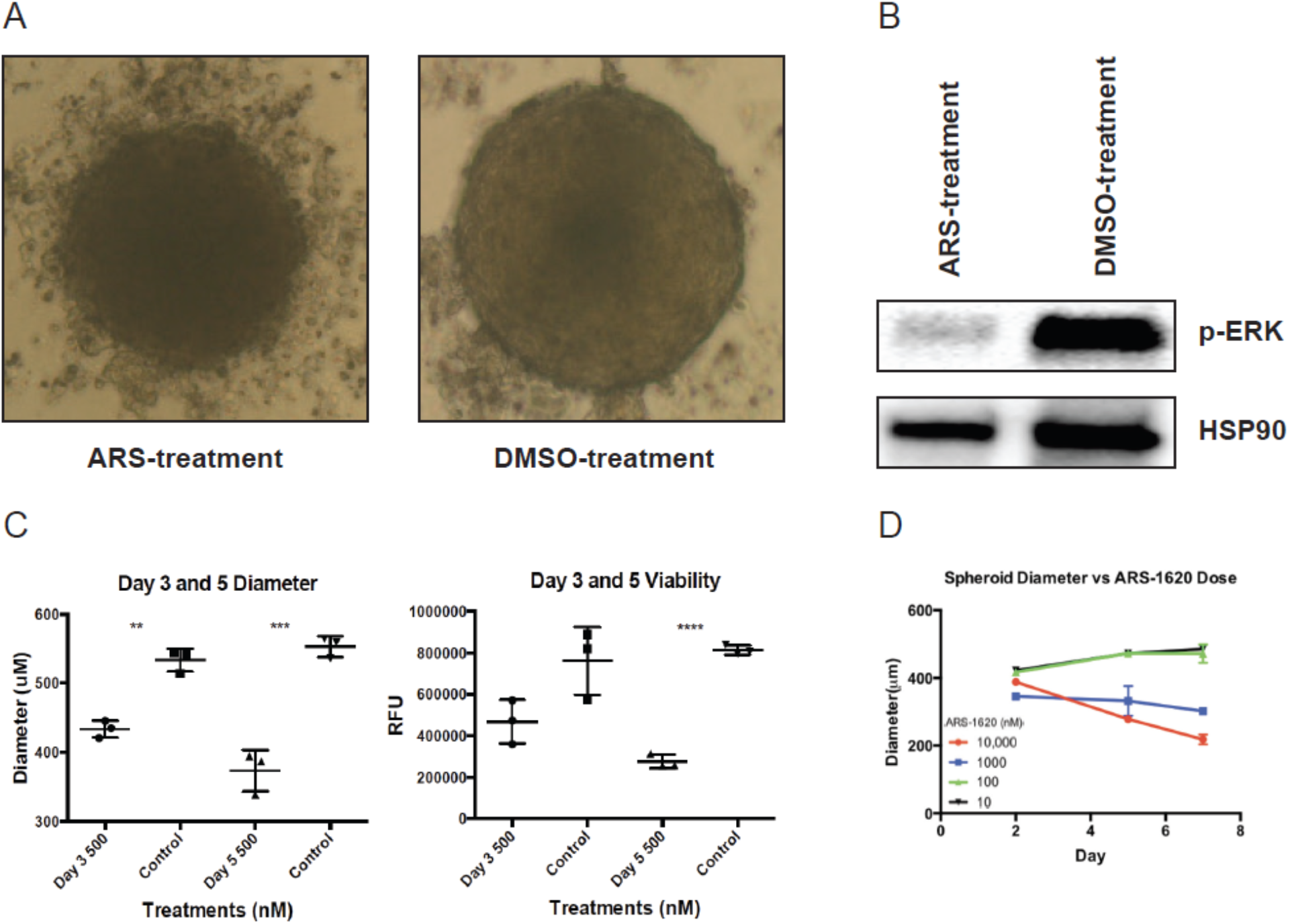
**A**. H358 3D lung cancer spheroids treated with ARS or DMSO. **B**. Western blot for p-ERK and HSP90 using H358 3D lung cancer spheroids treated with ARS or DMSO. **C**. Diameter measurements (in micrometers) (left plot) and cell viability (Cell Titer-Glo® luminescent cell viability in relative fluorescence units) (right plot) of H358 3D lung cancer spheroids treated with ARS or DMSO after 3 or 5 days of treatment (500 nM ARS-1620 or DMSO). **D**. Diameter measurements (in micrometers) of H358 3D lung cancer spheroids treated with different concentrations of ARS-1620 (nM) for 7 days.

## Notes

### Competing Interest Statement

The authors have declared no competing interest.

